# Long-Read Sequencing Reveals Increased Isoform Diversity in Key Transcription Factor Effectors of Intercellular Signalling at the Invertebrate-Vertebrate Transition

**DOI:** 10.1101/2024.11.12.623262

**Authors:** Nuria P. Torres-Aquila, Marika Salonna, Sebastian Shimeld, Stefan Hoppler, David E.K. Ferrier

**Affiliations:** The Scottish Oceans Institute, School of Biology, University of St Andrews, St Andrews, Fife, KY16 8LB, United Kingdom; Departament de Genètica, Microbiologia i Estadística, Facultat de Biologia, Institut de Recerca de la Biodiversitat (IRBio), Universitat de Barcelona, Barcelona, 08028, Spain; Institute of Medical Sciences, Foresterhill Health Campus, University of Aberdeen, Aberdeen, AB25 2ZD, United Kingdom; Institute of Education in Healthcare and Medical Sciences, Foresterhill Health Campus, University of Aberdeen, Aberdeen, AB25 2ZD, United Kingdom; The Department of Biology, University of Oxford, 11A Mansfield Road, Oxford, OX1 3SZ, United Kingdom

**Keywords:** TCF, SMADs, GLIs, *Ciona*, lamprey, *Lampetra planeri*, *Xenopus tropicalis*

## Abstract

Several intercellular signalling pathways (namely wingless - Wnt, Hedgehog - Hh, and Bone Morphogenetic Protein - BMP) are used repeatedly in animals throughout development and evolution, and are also frequent targets for disease-associated disruptions. We have previously shown that the major transcriptional effectors of β-catenin-dependent Wnt signalling, the TCF/LEF proteins, in contrast to other pathway components, have a higher gene number and isoform diversity in vertebrates versus invertebrates, but this increased diversity has only been poorly quantified. Considering that isoform diversity correlates with organism complexity, any increase in major signalling effectors is likely to have made a significant contribution to vertebrate evolution. Using *de novo* long-read transcriptomes, we compared isoform number per gene for the chordates *Ciona intestinalis, Lampetra planeri* and *Xenopus tropicalis*, thus encompassing the invertebrate sister group to vertebrates, as well as a cyclostome and a gnathostome vertebrate. Our results implicate an increase in isoform diversity of the transcription factors of major intercellular signalling pathways as having a disproportionate role in the evolutionary origin and diversification of vertebrates.

## Introduction

The driving forces for the evolution of organism complexity has been a topic of discussion for decades^1–4^. Despite genome duplications being renowned for the creation of new paralogous gene copies and their subsequent evolution via processes like sub- and neofunctionalization^5^, and specialisation^6^, the G-value paradox showed that the number of genes in a genome do not necessarily correlate with organism complexity^2^. One of the proposed alternatives to solve this paradox is the expansion of the organism proteome through alternative splicing, correlating with phenotypic novelty^3^. Previous studies found a strong correlation between number of cell types (as a proxy for organism complexity) and alternative splicing^4,7^, providing evidence of the importance of isoform diversity for organism evolution. However, whether particular types of genes contribute disproportionately to this phenomenon has not been assessed.

Wnt signalling is a cell-to-cell signalling mechanism highly conserved in the animal kingdom, required during development and regeneration^8^. The best described Wnt pathway is the canonical Wnt (cWnt) pathway, also known as the Wnt/β-catenin pathway, which involves the nuclear translocation of β-catenin, triggered by extracellular Wnt ligand-receptor interactions. Nuclear β-catenin functions as co-regulator for activation of Wnt-target genes usually via binding the T-cell factor/lymphoid enhancer factor (TCF/LEF) proteins. The cWnt pathway has a variety of roles in animal homeostasis and development, including involvement in development of the anterior-posterior and dorsal-ventral axes^8,9^. It is also associated in many human diseases such as cancers^10^, diabetes^11^ and mental disorders^12^. Comparably widespread functions in development, homeostasis and disease are also seen in other major signalling systems such as the Hedgehog (Hh) and Bone Morphogenetic Protein (BMP) pathways, whose main transcription factors are the Glioma-Associated Oncogene (GLI) proteins and the small/Mothers Against DPP Homolog (SMAD) proteins respectively^13–15^.

Genome comparisons between vertebrates and invertebrates reveal a remarkable conservation of the cWnt pathway with relatively little expansion of most of its components^16^. Nonetheless, vertebrate TCF/LEF transcription factors, the main transcription factor of the cWnt pathway, show a much greater diversity^17–20^. Multiple copies of *TCF/LEF* genes have been retained from genome duplications in vertebrates, which typically possess four TCF/LEF family genes with multiple isoforms, while invertebrates typically have one *TCF* gene with a single isoform^20,21^. A similar gene expansion might have occurred for SMAD and GLI families of TFs mediating BMP/TGFβ and hh signalling, respectively.

Thus, given these general observations we aimed to accurately assess the transcript isoform diversity across components of these signalling pathways and compare them to other categories of genes. We hypothesized that such major developmental control genes may have been a particular target for the evolutionary diversification that occurred with the origin of the vertebrates. We selected three species representing key lineages of the Olfactores chordates; the invertebrate urochordate *Ciona intestinalis*, the cyclostome (jawless vertebrate) *Lampetra planeri*, and a gnathostome (jawed vertebrate) *Xenopus tropicalis*, to analyse in an unbiased way the number of genes and transcripts expressed during embryogenesis and asses if TCF/LEF, SMAD and GLI genes are in some way distinctive in their transcript isoform diversity.

## Results

### Transcriptome analysis

To analyse the diversity of isoforms within chordates, we performed cDNA long-read sequencing of selected developmental stages of *C. intestinalis, L. planeri* and *X. tropicalis* (see Table 1) and processed the data following the pipeline shown Figure 1.

**Table 1.**
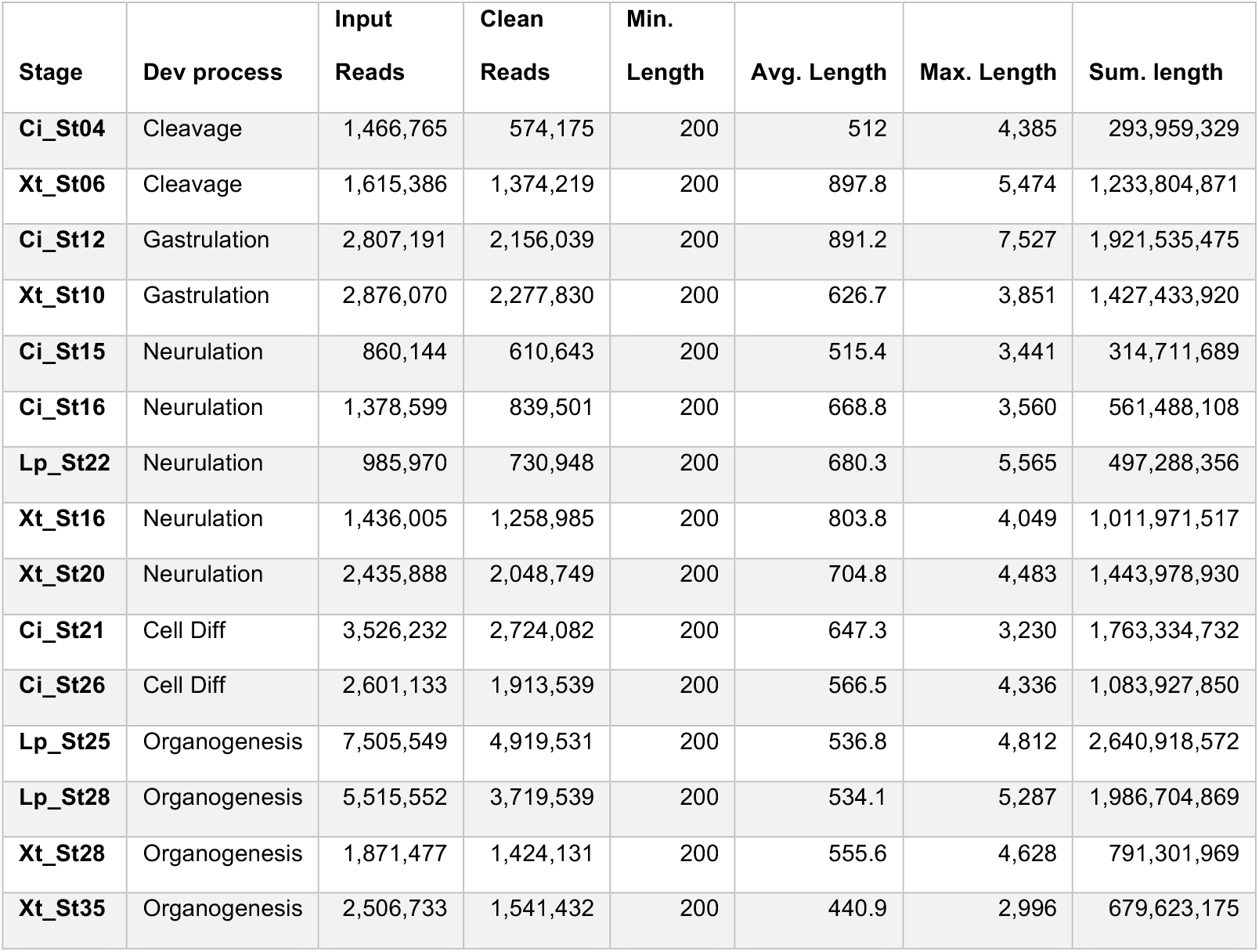
Long-read sequencing reads. Reads obtained after long-read sequencing and after data processing for each sample used. Ci: *Ciona intestinalis*; Lp: *Lampetra planeri*; Xt: *Xenopus tropicalis*; St: developmental stage.

| Stage | Dev process | Input<br>Reads | Clean<br>Reads | Min.<br>Length | Avg. Length | Max. Length | Sum. length |
| --- | --- | --- | --- | --- | --- | --- | --- |
| Ci_St04 | Cleavage | 1,466,765 | 574,175 | 200 | 512 | 4,385 | 293,959,329 |
| Xt_St06 | Cleavage | 1,615,386 | 1,374,219 | 200 | 897.8 | 5,474 | 1,233,804,871 |
| Ci_St12 | Gastrulation | 2,807,191 | 2,156,039 | 200 | 891.2 | 7,527 | 1,921,535,475 |
| Xt_St10 | Gastrulation | 2,876,070 | 2,277,830 | 200 | 626.7 | 3,851 | 1,427,433,920 |
| Ci_St15 | Neurulation | 860,144 | 610,643 | 200 | 515.4 | 3,441 | 314,711,689 |
| Ci_St16 | Neurulation | 1,378,599 | 839,501 | 200 | 668.8 | 3,560 | 561,488,108 |
| Lp_St22 | Neurulation | 985,970 | 730,948 | 200 | 680.3 | 5,565 | 497,288,356 |
| Xt_St16 | Neurulation | 1,436,005 | 1,258,985 | 200 | 803.8 | 4,049 | 1,011,971,517 |
| Xt_St20 | Neurulation | 2,435,888 | 2,048,749 | 200 | 704.8 | 4,483 | 1,443,978,930 |
| Ci_St21 | Cell Diff | 3,526,232 | 2,724,082 | 200 | 647.3 | 3,230 | 1,763,334,732 |
| Ci_St26 | Cell Diff | 2,601,133 | 1,913,539 | 200 | 566.5 | 4,336 | 1,083,927,850 |
| Lp_St25 | Organogenesis | 7,505,549 | 4,919,531 | 200 | 536.8 | 4,812 | 2,640,918,572 |
| Lp_St28 | Organogenesis | 5,515,552 | 3,719,539 | 200 | 534.1 | 5,287 | 1,986,704,869 |
| Xt_St28 | Organogenesis | 1,871,477 | 1,424,131 | 200 | 555.6 | 4,628 | 791,301,969 |
| Xt_St35 | Organogenesis | 2,506,733 | 1,541,432 | 200 | 440.9 | 2,996 | 679,623,175 |

**Figure 1.**
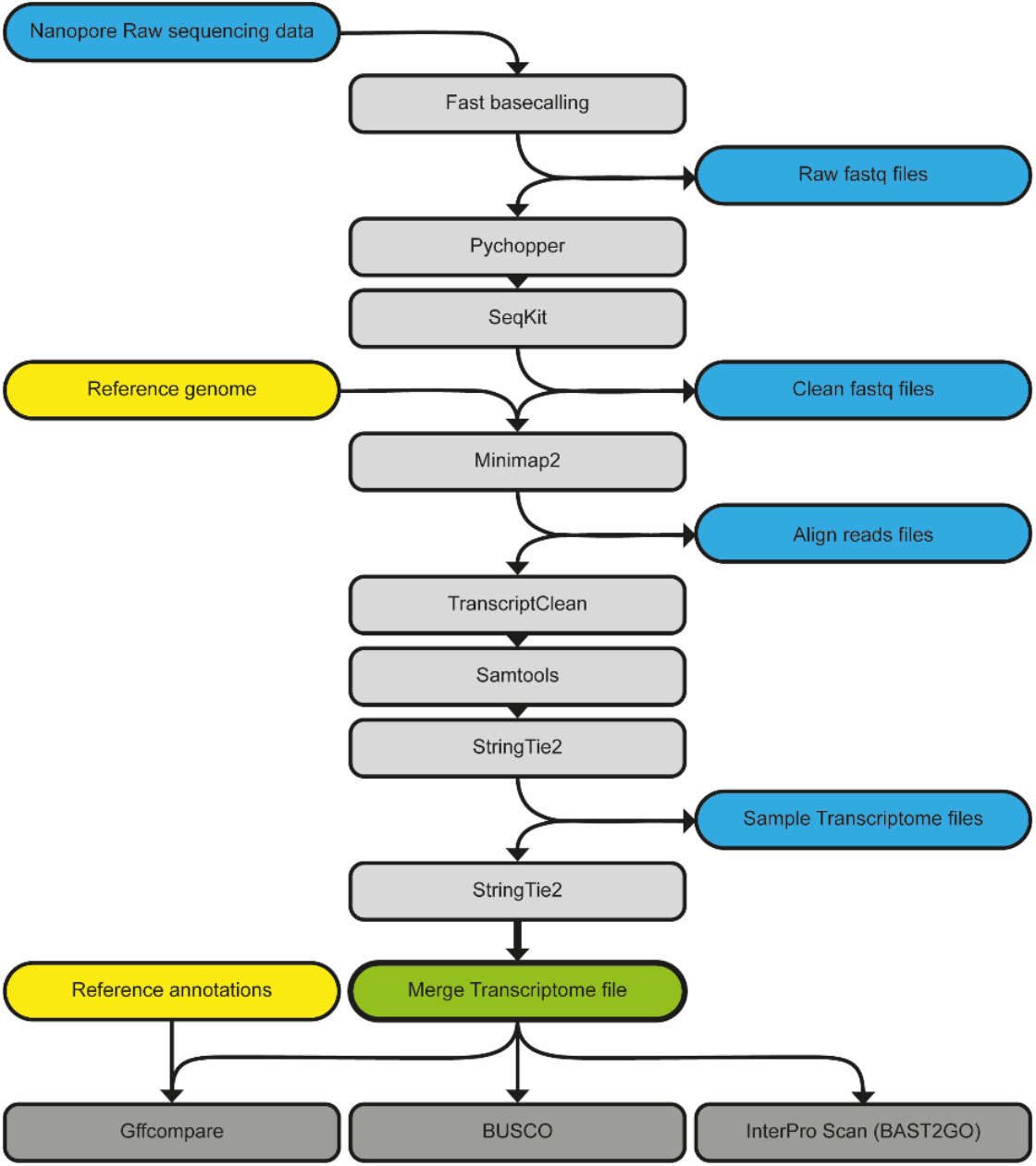
Pipeline for long-read data processing. Round boxes: files; Square boxes: software programs. Blue boxes: obtained files; Yellow boxes: reference files; Green box: final transcriptome file; Light-grey boxes: software for processing data; Dark-grey boxes: software for evaluate quality of obtained transcriptome file.

All selected stages performed similarly in the sequencing protocol (Table 1) producing *de novo* transcriptomes with loci coverage over 40% (Figure 2A) and with over 60% of metazoan-conserved orthologs (BUSCOs) found (Figure 2D). Notably, the transcriptomes included over 10% of novel loci, with gene models not currently annotated in the respective reference genomes (Figure 2B), besides the highest proportion of transcripts being ones that fully match the reference models (categories ‘=‘, ‘c’ and ‘k’ of Figure 2C). Regarding the novel loci, the vast majority of the identified genes had no GO term associated with them (1110 out of 1359 for *C. intestinalis*, 3184 out of 3337 for *L. planeri*, and 2898 out of 3316 for *X. tropicalis*), suggesting they may be taxon-specific or rapidly evolving genes. After performing a GO enrichment analysis on the loci that did have associated GO terms, no GO terms were enriched for *C. intestinalis* or *L. planeri*. However, for *X. tropicalis* we found 209 GO terms significantly enriched (Supplemental Table 1), most of them linked to muscle- and heart-related functions, including muscle contraction, structural assembly, and development.

**Figure 2.**
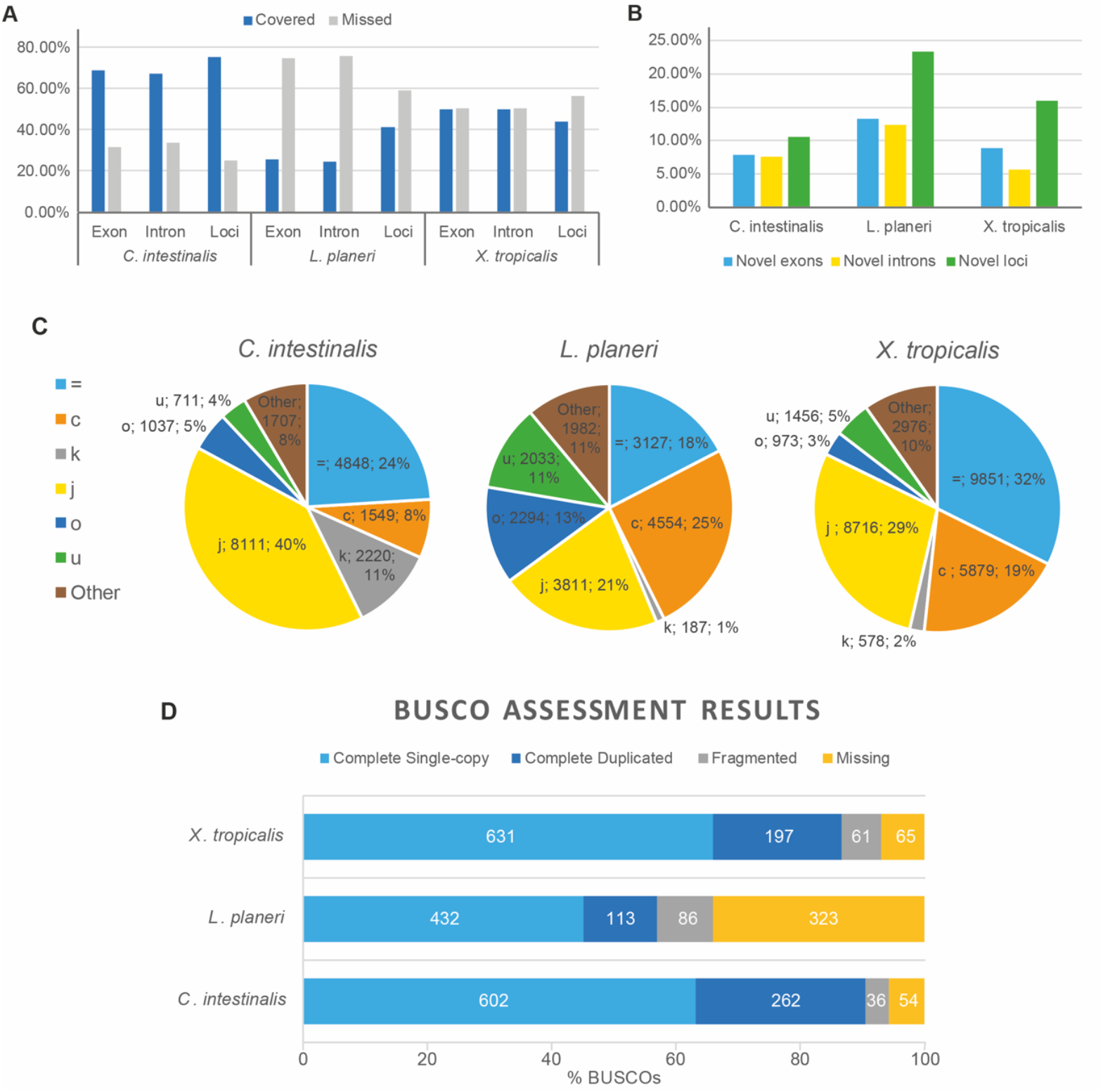
Transcriptome quality assessments. A) Covered (blue) and missed (grey) exons/intron/loci for each species data set. B) Novel exon (blue), intron (yellow), and loci (green) for each specie data set. C) Pie plots showing the distribution of transcript-matching types according to gffcompare categories: = (light-blue), identical reference-query match; c (orange), complete query match within reference; k (grey), complete reference match within query; j (yellow), some splice site mismatch; o (dark-blue), partial overlapping match; u (green), no match of query within reference (novel); Other (brown), other types of query-reference matches including m (all introns retain), n (some introns retain), e (single exon match), s (intron match on opposite strand), x (exon match on opposite strand), i (contained within reference intron), y (reference contained within intron), p (possible polymerase run-on), and r (repeat). D) Histogram showing the amount of evolutionary conserved orthologs (BUSCOs) found, as single-copy (light blue), duplicated (dark blue) or fragmented (grey), and missing (yellow).

The transcript:gene ratio (t/g ratio) was calculated for the total number of genes and transcripts obtained for each transcriptome, as well as for subsets of particular categories of genes (Table **2**). We observed higher number of transcripts per gene in the vertebrates relative to invertebrates only for TCF/LEF genes (TCFs), SMADs and GLIs. Comparisons of the variance of t/g ratio observed on each subset showed that TCF/LEF genes (TCFs) had a greater variance than the other gene subsets studied (p-value < 0.1, p_A_ values of Figure 3), including the subset “DNA-binding Transcription Factor activity” (GO:0003700) and the SOX genes, which belong to the same HMG-box superfamily as the TCF/LEFs. Moreover, the same results were found for SMAD and GLI genes with statistically significant support (p-value < 0.05, SMADs and GLIs, p_B_ and p_C_ values of Figure 3, respectively). This indicates that these key developmental transcription factors (TCF/LEFs, SMADs, and GLIs) have a distinctive pattern of a higher number of transcripts per gene in the vertebrates relative to invertebrates: in our analysis, TCF/LEFs have 1 gene and 1 transcript in *C. intestinalis* compared to 1 gene and 2 transcripts in *L. planeri* and 4 genes and 9 transcripts in *X. tropicalis*; SMADs have 2 genes and 2 transcripts in *C. intestinalis* compared to 4 genes and 7 transcripts in *L. planeri* and 6 genes and 17 transcripts in *X. tropicalis*; and GLIs have 1 gene and 1 transcript in *C. intestinalis* compared to 2 genes and 3 transcripts in *L. planeri* and 2 genes and 6 transcripts in *X. tropicalis*. Interestingly, the variance observed in the t/g ratio for these three gene families (TCF/LEFs, SMADs, and GLIs) was not significantly different between each of them (SMADs-TCFs, p-value = 0.34; GLIs-TCFs, p-value = 0.29; GLIs-SMADs, p-value = 0.44), indicating that they show similar distributions of transcripts and genes within the three chordate groups.

**Table 2.** Total number of genes and transcripts for each transcriptome and subsets. All: the whole transcriptome; Over 1: only genes with more than one transcript; Emb. Dev.: genes with the GO term ‘Embryo Development’ (GO:0009790); Wnt Path.: genes with the GO term ‘Wnt Pathway’ (GO:0016055); TF: genes with the GO term ‘DNA-binding transcription factor activity’ (GO:0003700); TCFs: *TCF* and *TCF/LEF* genes; SMADs: *SMAD* genes; GLIs: *GLI* genes; SOXs: *SOX* genes.

|  | *Ciona intestinalis* |  |  | *Lampetra planeri* |  |  | *Xenopus tropicalis* |  |  |
| --- | --- | --- | --- | --- | --- | --- | --- | --- | --- |
|  | Genes | Transcripts | Ratio t/g | Genes | Transcripts | Ratio t/g | Genes | Transcripts | Ratio t/g |
| All | 12,717 | 20,183 | 1.59 | 14,311 | 17,988 | 1.26 | 20,775 | 30,429 | 1.46 |
| Over 1 | 3,868 | 11,171 | 2.89 | 2,235 | 5,860 | 2.62 | 5,431 | 15,105 | 2.78 |
| Emb. Dev. | 15 | 22 | 1.47 | 14 | 19 | 1.36 | 69 | 106 | 1.54 |
| Wnt Path. | 40 | 58 | 1.45 | 29 | 33 | 1.14 | 89 | 134 | 1.51 |
| TF | 296 | 481 | 1.63 | 130 | 167 | 1.28 | 750 | 1235 | 1.65 |
| TCFs | 1 | 1 | 1.00 | 1 | 2 | 2.00 | 4 | 9 | 2.25 |
| SMADs | 2 | 2 | 1.00 | 4 | 7 | 1.75 | 6 | 17 | 2.83 |
| GLIs | 1 | 1 | 1.00 | 2 | 3 | 1.50 | 2 | 6 | 3.00 |
| SOXs | 5 | 7 | 1.40 | 3 | 5 | 1.67 | 9 | 14 | 1.56 |

**Figure 3.**
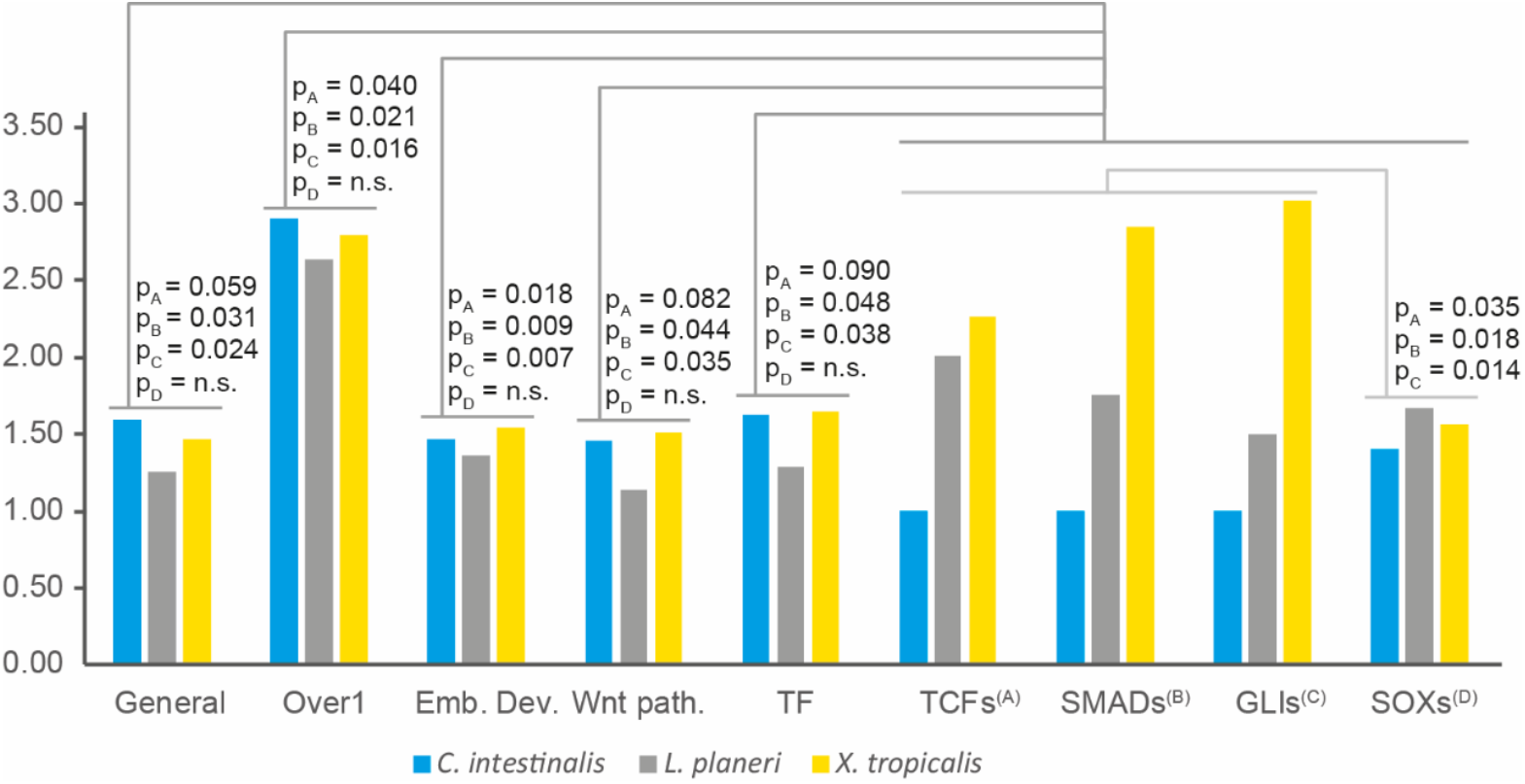
Ratio t/g. Histogram of the transcript:gene ratios (t/g ratio) for each species (blue: *Ciona intestinalis*; gray: *Lampetra planeri*; yellow: *Xenopus tropicalis*) and category (All: whole transcriptome; Over 1: genes with more than one transcript; Emb. Dev: genes with the GO term ‘Embryo Development’ (GO:0009790); Wnt path.: genes with the GO term ‘Wnt Pathway’ (GO:0016055); TF: genes with the GO term ‘DNA-binding transcription factor activity’ (GO:0003700); TCFs: *TCF* and *TCF/LEF* genes; SMADs: *SMAD* genes; GLIs: *GLI* genes; SOXs: *SOX* genes). p: F-test p-value, subscript indicates the gene family of the comparison subset corresponding to the superscript on the group name (A: TCFs, B: SMADs, C: GLIs; D: SOXs).

### A new splice isoform in *Ciona intestinalis* Type B

To confirm the transcript sequence and structure of the *C. intestinalis* TCF/LEF (in what was formerly known as *Ciona intestinalis* Type B) relative to the commonly studied sister species *Ciona robusta* (formerly *Ciona intestinalis* type A), we performed RACE-PCR in different stages of development, selected according to the previously described expression of *C. intestinalis TCF* (*CiTCF*)^22^. 5’RACE-PCR was performed in St04 (8-cell; maternal mRNA) and St12 (mid Gastrula; zygotic mRNA), and 3’RACE-PCR was performed for St04, St12, St16 (late Neurula), and St21 (mid Tailbud I).

For 5’RACE-PCR, only one fragment was amplified, matching the described gene model. For 3’RACE-PCR two different 3’ ends were found. The first was found in all the assessed stages and matched the previously described gene model. The second, smaller in size, was found in St16 and matched the described gene model but with a different final exon. Further analysis of the genomic region between exon 12 and exon 13 in *C. intestinalis* (intron 12; Figure 4A) showed the presence of this new exon flanked by two transposable elements, partially overlapping one at the 3’ end. These transposable elements matched in sequence the previously described miniature inverted-repeat transposable elements (MITE) Cimi-1^23^. Comparison of intron 12 between *C. robusta* and *C. intestinalis* revealed that despite the Cimi-1 insertions being conserved, this was not the case for the splice acceptor site nor the stop codon of exon 12.5 (Figure 4B), indicating that this newly discovered exon may be specific to *C. intestinalis*. To confirm this alternative C-terminus, we designed a specific Reverse primer for exon 12.5 and performed RT-PCRs on St12, St15 (mid Neurula) and St21. This isoform was found only in post-gastrulation stages. *C. intestinalis* thus produces two isoforms of TCF/LEF, in contrast to the single isoform produced by this gene in *C. robusta*.

**Figure 4.**
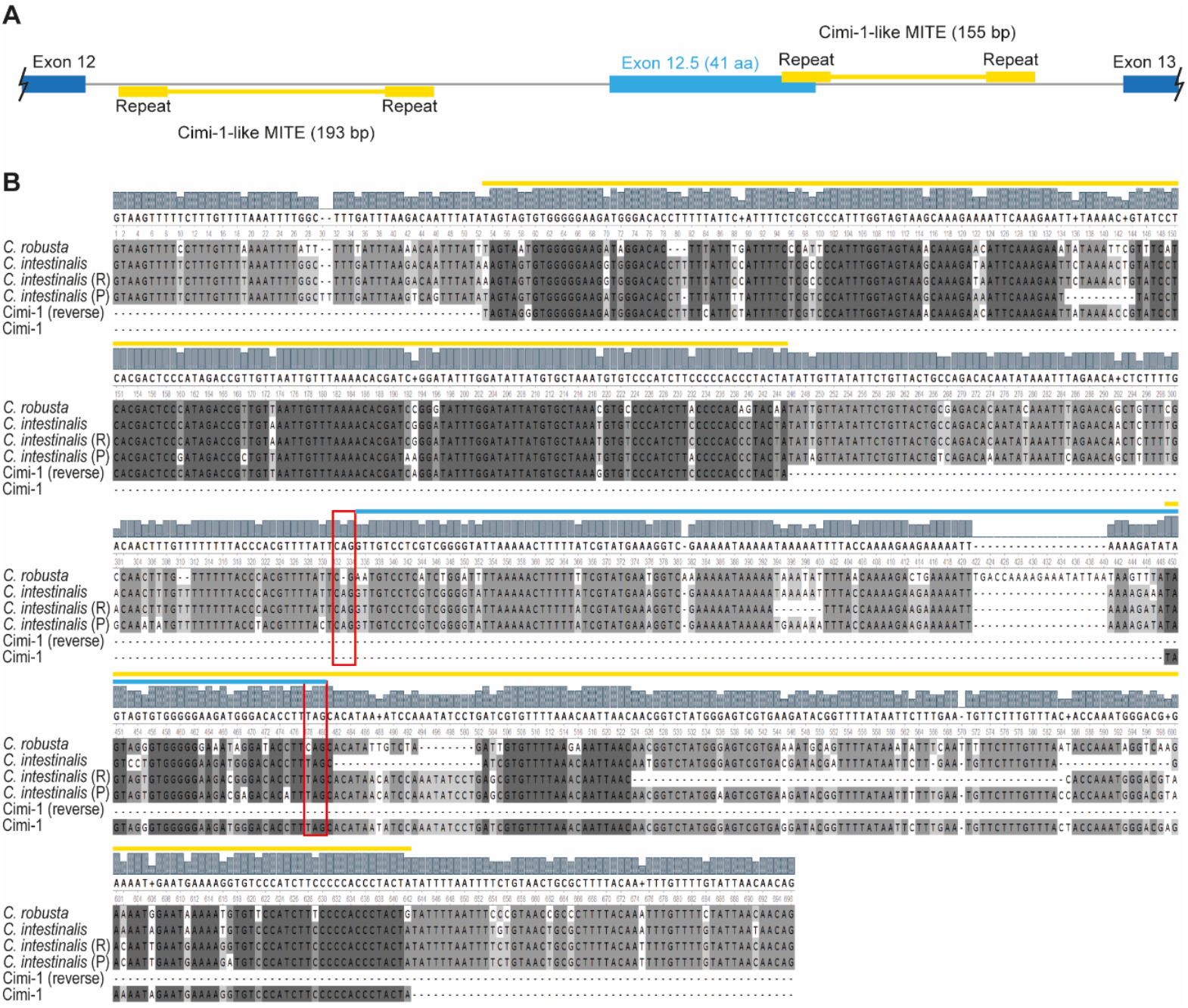
Schematic of *CiTCF* intron 12. A) Schematic of the *C. intestinalis* genomic region where exon 12.5 is found. B) DNA alignment of TCF intron 12 of *C. intestinalis* and *C. robusta. C. intestinalis* (R): sequence from Roscoff reference genome (GCA_018327825.1); *C. intestinalis* (P): sequence from Plymouth reference genome (GCA_018327805.1); dark blue: annotated exons; light blue: new exon (exon 12.5); yellow: Cimi-1-like sequences; red boxes: acceptor site and stop codon of exon 12.5.

## Discussion

Our transcriptome analyses focused on gene expression during embryo development and organogenesis revealed that *TCF/LEF, SMAD* and *GLI* genes exhibit a distinctive pattern of higher numbers of splice isoforms per gene in vertebrates than the representative from the closest invertebrate sister group, represented here by the urochordate *C. intestinalis*. This larger number of splice isoforms occurs in addition to the increase in paralog numbers in these transcription factors at the origin of vertebrates.

Our previous *in silico* analyses demonstrated that there was an increase in TCF/LEF gene number at the invertebrate-to-vertebrate transition, likely via the whole genome duplications that occurred early in vertebrate evolution, and in addition there was a dramatic increase in isoform diversity in this gene family^24^. However, this diversity remained to be quantified, as well as being related to when these isoforms can be detected during development and hence are presumably functional. This more detailed analysis is provided here. In addition, another open question was whether the evolutionary patterns seen in this major transcriptional effector of the cWnt signalling pathway was unique or was also observed elsewhere, such as in other major signalling pathways like hh and BMP.

Invertebrates have four SMADs, corresponding to a single-copy of each SMAD subgrouping (common SMAD (Co-SMAD), inhibitory SMAD (I-SMAD), BMP-regulated receptor SMAD (R-SMAD) and TGFβ-regulated R-SMAD), while vertebrates present eight different SMADs (one Co-SMAD, two I-SMADs, three BMP-regulated R-SMADs, and 2 TGFβ-regulated R-SMADs)^25,26^. In the GLI family, invertebrates have a single *GLI* gene while vertebrates typically have three paralogues^27^. Thus, the SMAD and GLI families show a similar pattern of increased paralog number to that seen in the TCF/LEF family. However, the amount of splice isoform diversity remained to be analysed for the SMAD and GLI families. Here we demonstrate that these families show a similar pattern to the TCF/LEF family, with a significantly higher number of isoforms per gene in vertebrates (here represented by a lamprey, *L. planeri*, and frog, *X. tropicalis*) relative to the invertebrate sister group (represented by the urochordate *C. intestinalis*).

It is striking that this invertebrate:vertebrate pattern for the TCF/LEF, SMAD and GLI genes is significantly different from all genes and their transcripts found in our new transcriptomes (‘General’ category in Fig.3). This is also the case when the focus is more specifically on those genes that exhibit alternative splicing and have more than one transcript per gene (‘Over 1’ category in Fig.3), in which the *C. intestinalis* ratio is indistinguishable from those of the two vertebrates. These first two categories encompass genes that span a variety of biological functions and so we also focused on genes thought to be more specifically involved in embryo development, in case there is a general increase in complexity of developmental control genes associated with the invertebrate-to-vertebrate transition and the evolution of vertebrate complexity. No distinct pattern was observed between the invertebrate *C. intestinalis* and the vertebrates. This lack of invertebrate to vertebrate distinction was also observed when the focus was even more specific, onto Wnt pathways as a whole (Fig.3). Another alternative possibility was explored by comparison to Transcription Factors in general (‘TF’ in Fig.3) in case the mediators of transcriptional control are the focus of change between invertebrates and vertebrates. No significant distinction was found. As a final test of how distinct the pattern found for the TCF/LEF family was, we analysed the SOX genes, since these are in the same superfamily as the TCF/LEF genes and hence act as the closest comparison possible. The invertebrate:vertebrate pattern for the TCF/LEF genes is significantly different to that of the SOX genes. Thus, the TCF/LEF genes stand-out from all of these different categories of genes, implying a specific expansion in the splice isoform diversity focused on these transcription factor mediators of the cWnt pathway. Notably, the only categories of genes that we found with comparable invertebrate:vertebrate patterns were those of the transcription factor mediators of other major intercellular signalling pathways (the SMAD and GLI genes).

Amongst all of this vertebrate genetic diversity, it has been shown that vertebrate TCF/LEF paralogs and GLI paralogs have some degree of redundancy at the functional level^17,28^. Nevertheless, there is also evidence that different TCF/LEF isoforms can target different genes ^29^, showing sub/neo-functionalization. Similarly, GLI isoforms have been shown to have opposing roles activating or repressing the gene expression of their specific target genes^28^. Therefore, the diversity of vertebrate genes in TCF/LEF, SMAD and GLI families, and the isoforms produced from them, presumably reflects a wide array of functional capabilities downstream of important developmental signalling pathways in vertebrates.

These three gene families are the main transcription factor effectors of major intercellular signalling pathways (cWnt, BMP/TGFβ and Hh, respectively), which are integral to embryo development, organogenesis and homeostasis. Their major roles in development in conjunction with the correlation of isoform diversity with organism complexity^1–4^ is consistent with the hypothesis that increased diversity of these transcription factors may be making a disproportionate contribution to the evolution of vertebrate complexity relative to invertebrates. Interestingly, previous studies had observed genes resulting from duplication usually retain lower numbers of isoforms, with duplication and isoform diversity being inversely correlated evolutionary mechanisms^30,31^. This makes the pattern observed here in these developmental signalling transcription factor effectors even more striking, as our data shows that *TCF/LEF, SMAD* and *GLI* genes are exceptions to this inverse correlation. This could be an indicator of a significant role for isoform diversity of these key transcription factors in the evolutionary origins and diversification of vertebrate complexity.

One caveat to this hypothesis is whether the species selected here are good representatives of the invertebrate-to-vertebrate transition. *C. intestinalis*, for example, was selected because it is a urochordate and as such is a member of the closest invertebrate clade to the vertebrates, and is also accessible and amenable to gene expression and developmental experimentation. There are, however, aspects of its genome organisation and content that are relatively derived within the chordates^32^. Also, it is known that amphioxus exhibits alternative splicing from its *Gli* gene, producing two distinct isoforms^33^, rather than the single isoform of *Ciona* GLI that was found here. It is notable, however, that our analyses are focused on the transcripts found within our new transcriptome data and are not necessarily capturing all isoforms produced by each of the three species selected. Rare transcripts expressed at very low levels in embryogenesis, or transcripts expressed only in adult stages, will not be present in our data. These are areas for future further work, to quantify the patterns described here with even greater precision. Nevertheless, there is no indication that the three transcription factor families focused on here are unusual in *Ciona* relative to invertebrates in general in any major way.

In addition to these “signalling transcription factor” findings, the long-read transcriptomes provided in this work are also a valuable resource for deeper understanding of gene expression during embryo development of different chordates, including species not previously assessed with long-read transcriptome sequencing, such as Cyclostomata and Urochordata. However, despite all the samples being processed in the same way and our obtaining similar quality values within urochordate and gnathostome data sets, the quality of the cyclostome data set was not as good (Figure 2). This issue could be due to the GC-richness of cyclostome genomes^34^ and/or the fact that *L. planeri* (the species sampled) and *P. marinus* (the species used as the lamprey reference genome) are more evolutionary distant and distinct than the species used for urochordates (samples of *C. intestinalis* and reference genome of *C. robusta*) and gnathostomes (samples and reference genome of *X. tropicalis*).

It is also notable that for *C. intestinalis*, the transcript type category that had the highest number of transcripts was ‘j’ (40%, Figure 2C), showing that most of the transcripts had mismatched splicing sites (junctions) against reference annotations. This could be an indicator of potential novel isoforms found in this new *C. intestinalis* transcriptome data for previously annotated genes in *C. robusta*. In fact, our finding of an alternative transcript of *CiTCF* is the first evidence that *C. intestinalis* has more than one TCF isoform. However, this second transcript of *CiTCF* was only found by RACE-PCR rather than being in the transcriptome data, which may reflect the RACE-PCR having higher sensitivity than cDNA long-molecule sequencing, since the PCR is gene-specific. In addition, the partial overlap of exon 12.5 with the Cimi-1 MITE sequence provides an example of how transposable elements can alter intronic sequences that then provide material for the evolutionary origin of novel exons, in this case in concert with point mutations that generated a new splice acceptor site as well as a new stop codon.

In conclusion, we have created *de novo* transcriptomes of embryo development for three different chordates: *C. intestinalis* (Urochordata), *L. planeri* (Cyclostomata), and *X. tropicalis* (Gnathostomata). Our analyses demonstrate distinctive increases in isoform diversity at the invertebrate-to-vertebrate transition specifically among transcription factor effectors of key intercellular signalling pathways that drive cell type diversity. This distinctive change focused on these specific gene families (TCF/LEF, SMAD, and GLI) goes beyond the previous observations of a general correlation between increased isoform diversity and evolution of animal complexity. This demonstrates likely disproportionate roles for these specific transcription factor families in the evolution of vertebrate complexity, which needs to be explored with future functional assays of these various isoforms.

## Methods

### Material fixation and RNA extraction

After *in vitro* fertilization, selected embryological stages from *C. intestinalis, L. planeri* and *X. tropicalis* were fixed in RNA*later*™ (Invitrogen, AM7021) for a minimum of 16h at 4°C, taking care that the amount of RNA*later*™ was at least 10 times the volume of the sample. Stage numbering was done accordingly to Hotta^35^, Tahara^36^, and Zahn^37^. RNA extractions were performed with ‘RNAeasy mini kit’ (QIAGEN, 74104) following the manufacturer’s protocol. The quality and quantity of the total RNA obtained was tested by gel electrophoresis and Nanodrop spectrophotometer.

### cDNA long-read sequencing

For each sample analysed, 50ng of total RNA was processed using the PCR-cDNA Barcoding Kit (Oxford Nanopore Technologies (ONT), SQK-PCB109). The first strand synthesis was performed following the manufacturer’s instructions (Thermo Scientific Maxima H Minus First Strand cDNA Synthesis Kit with dsDNase, K1681). First, a previous DNAse treatment of the total RNA was performed as follows: incubation of 10 min at 37°C followed by an inactivation of 5 min at 55°C in the presence of 10 mM of DTT. After, the sample was cooled on ice and VN Primers and dNTPs where added. After mixing, the sample was incubated 5 min at 65°C and snap cooled on a pre-chilled freezer block. Then, 5xRT buffer, Strand-Switching Primers and Maxima H Minus Enzyme Mix were added and the sample was mixed by pipetting. The reverse transcription (RT) reaction was performed by incubating the sample 10 min at 25°C followed by 92 min at 42°C and 5 min at 85°C.

A single PCR reaction was performed for each RT reaction. The PCR reaction was prepared according to the PCR-cDNA Barcoding Kit protocol with minor modifications in the cycling conditions: initial denaturalization at 95°C for 30s; 18 cycles of denaturation at 95 °C for 10s, annealing at 62 °C for 20s, extension at 65 °C for 2min 30s; final extension of 65 °C for 10 min and hold at 4 °C. Each reaction was treated with Exonuclease I (New England Biolabs, M0293) followed by a purification with AMPure XP beads (Beckman Coulter, A63880) as indicated in the PCR-cDNA Barcoding Kit protocol with minor modification (i.e. we used 30µL of AMPure XP beads per PCR reaction). The concentration and quality of the obtained samples were assayed by Nanodrop and gel electrophoresis. The sequencing was performed with a maximum of 100 fmol per run.

MinION flow cells (ONT, FLO-MIN106D) underwent flow cell check prior to library construction. The barcoded PCR-cDNA libraries were prepared for sequencing and the MinION flow cell was primed using the flow cell priming kit (ONT, EXP-FLP002) as indicated in the PCR-cDNA Barcoding Kit protocol. A maximum of six PCR-cDNA libraries per run were sequenced in parallel on a single MinION flow cell with Min-KNOWN software v.21.02.2 (ONT). Fast basecalling was performed in real-time with a maximum data acquisition time of 48h and the following filters applied: minimum Barcode score of 60 and minimum Qscore of 7. All the raw data produced is available at the Sequence Read Archive (SRA) database under accession numbers SRR24756885-SRR24756899, BioProject PRJNA977127.

### Transcriptomic data processing

The cDNA ONT reads were pre-processed with pychopper (ONT), to remove the primer sequences introduced by the protocol, and with Seqkit^38^, to remove sequences under 200bp length. Each cDNA ONT library for each developmental stage sequenced was aligned to the corresponding reference genome by minimap2^39^ (*C. intestinalis* reads aligned against *Ciona robusta* genome (GCA_000224145.2); *L. planeri* reads aligned against *Petromyzon marinus* genome (GCA_010993605.1); *X. tropicalis* reads aligned against *X. tropicalis* genome (GCA_000004195.3)), transcripts refined with TranscriptClean^40^ and annotated with StringTie2^41^ software. Finally, the stage-specific annotations were merged into a general annotation file for each species with StringTie2 *‘-merge’* option and reference transcriptome and proteome dataset were generated using TransDecoder^42^. The obtained proteome datasets and general annotation file were analysed with BUSCO^43^ (v5.2.2) and gffcompare^44^, respectively, to assess the quality of the transcriptomes. For obtaining the GO annotations, the InterPro Scan option of Blast2GO software (v.6.0.3) and the EggNOG-mapper tool^45^ were run using each transcriptome dataset as input.

### Statistical analysis

The GO enrichment analysis was done separately for each species. The analysis was performed in R using the function ‘enricher()’ from the ‘clusterProfiler’ package providing as inputs the list of ‘u’ genes and the GOs and genes found in the whole transcriptome.

For each species dataset, the ratio transcript:gene (t/g ratio) was calculated for all the genes present in the created transcriptome (All), genes that had more than 1 transcript (Over 1), genes with the gene ontology number GO:0009790 (Embryo Development, Emb. Dev.), genes with the gene ontology number GO:0016055 (Wnt pathway, Wnt Path.), genes with the gene ontology number GO:0003700 (DNA-binding transcription factor activity, TF) and for the gene families TCFs, SMADs and GLIs. We performed F-tests to compare the variances of t/g ratios within the different subsets (i.e. All, Over 1, Emb. Dev., Wnt Path., TF, TCFs, SMADs, and GLIs) doing pairwise comparisons. A difference in the variance was expected when the t/g ratio was noticeably different within species.

### RACE-PCR, cloning and sequencing

Total RNA of *C. intestinalis* was used for 3’RACE and 5’RACE experiments using FirstChoice™ RLM-RACE Kit (Invitrogen, AM1700) following the manufacturer’s protocol and with the following CiTCF-specific primers: 5’-CAGGCATGTTACGATACCCATATCCA-3’ (3’RACE), 5’-

CATCACAATTTACATCCACATCTGGTGGT-3’ (5’RACE), 5’-CTTCACATATGGCCGACTTGGTTTGTCACCT-3’ (nested 3’RACE) and 5’-

TCGCGTTTCTTTGAACCAGGTTCAG-3’ (nested 5’RACE). PCRs were performed with Taq DNA polymerase (Thermo Scientific, EP0402) following the manufacturer’s protocol and results were assessed by gel electrophoresis in 1% agarose gels.

The individual bands obtained after nested RACE-PCR were purified with the ISOLATE II PCR and Gel Kit (Meridian Bioscience, BIO-52059), and cloned with the pGEM®-T Easy Vector System (Promega, A1360), following the manufacturer’s protocol. Transformation was performed into *E. coli* competent cells (Aligent, XL10-Gold Ultracompetent cells, 200314) by heat-shock. All clones were selected by Ampicillin resistance and their composition was confirmed by enzyme digestion with NotI (New England Biolabs, R0189) and Sanger sequencing (Oxford Zoology service, Eurofins service).

The intronic region within exon 12 and exon 13 of *C. intestinalis* TCF was amplified with the following primer pair: 5’-TCGCACGATAATGTTAACAAGC-3’ (forward primer), 5’-GTTCATAGCTACTTGATGGTTGGA-3’ (reverse primer). Specific exon 12.5 PCRs were done with the following primer pair: 5’-ACAACAGCAATTATGGTGCGCAC-3’ (forward primer), 5’-ATACCCCGACGAGGACAAC-3’ (reverse primer). The protocols for PCR and posterior cloning were as described above.

## Acknowledgments

We thank members of the Ferrier and Hoppler labs for discussions related to this work.

